# SpeciesGeoCoder: Fast categorisation of species occurrences for analyses of biodiversity, biogeography, ecology and evolution

**DOI:** 10.1101/009274

**Authors:** Mats Töpel, Maria Fernanda Calió, Alexander Zizka, Ruud Scharn, Daniele Silvestro, Alexandre Antonelli

## Abstract

Understanding the patterns and processes underlying the uneven distribution of biodiversity across space and time constitutes a major scientific challenge in evolutionary biology. With rapidly accumulating species occurrence data, there is an increasing need for making the process of coding species into operational units for biogeographic and evolutionary analyses faster, automated, transparent and reproducible. Here we present SpeciesGeoCoder, a free software package written in Python and R, that allows for easy coding of species into user-defined areas. These areas may be of any size and be purely geographical (i.e., polygons) such as political units, conservation areas, biomes, islands, biodiversity hotspots, and areas of endemism, but may also include altitudinal ranges. This flexibility allows scoring species into complex categories, such as those encountered in topographically and ecologically heterogeneous landscapes. In addition, SpeciesGeoCoder can be used to facilitate sorting and cleaning of occurrence data. The various outputs of SpeciesGeoCoder include quantitative biodiversity statistics, global and local distribution maps, and NEXUS files that can be directly used in many phylogeny-based applications for ancestral state reconstruction, investigations on biome evolution, and diversification rate analyses. Our simulations indicate that even datasets containing hundreds of millions of records can be analysed in relatively short time using a regular desktop computer. We exemplify the use of our program through two contrasting examples: *i)* inferring historical dispersal of birds across the Isthmus of Panama, separating lowland *vs.* montane species and optimising the results onto a species-level, dated phylogeny; and *ii)* exploring seasonal variations in the occurrence of 10 GPS-tracked individuals of moose (*Alces alces*) over one year in northern Sweden. These analyses show that SpeciesGeoCoder allows an easy, flexible and fast categorisation of species distribution data for various analyses in ecology and evolution, with potential use at different spatial, taxonomic and temporal scales.

## Introduction

Species distributions provide the basic knowledge for biodiversity research, allowing us to understand their environmental requirements, biogeographic history, and expected resilience to climate change. However, analysing the distribution of the world’s estimated 8.7 million species (Mora, C., Tittensor, D.P., et al. 2011) remains a major scientific challenge.

There are now over half a million species occurrences worldwide freely available through the Global Biodiversity Information Facility (GBIF; http://www.gbif.org) and their partners, of which the majority (about 446 million) are geo-referenced, i.e. provided with a latitude and longitude. These numbers are steadily increasing thanks to new agreements on data sharing, on-going digitalisation programmes, the work of active field biologists and amateurs combined with an increasing awareness of the need to provide rich metadata with biological collections and sequences (Hyde KD, U.D., Manamgoda DS, Tedersoo L, Nilsson RH. 2013), and tools that enable automated geo-referencing of older museum specimens (Guralnick, R.P., Wieczorek, J., et al. 2006, Garcia-Milagros, E. and Funk, V.A. 2010). Publicly available species occurrences represent an enormous data source for biodiversity research, but are as yet poorly exploited due to two main factors: *i)* general scepticism concerning the quality of records available, in terms of species identification and precise coordinates (Hjarding, A., Tolley, K.A., et al. 2014), and *ii)* demonstrated taxonomic, geographic, and temporal biases (Boakes, E.H., McGowan, P.J., et al. 2010). Whereas these biases are slowly being compensated by data growth, improving quality typically relies on expert curation and targeted evaluations (e.g. http://www.iucnredlist.org/) as well as tools for massive data cleaning (e.g. through workflows at the Biodiversity Virtual e-Laboratory, http://www.biovel.eu).

Research using species distribution data has been further hampered by a lack of tools that allow for classification of raw occurrences into discrete categories. This is often a crucial step in many evolutionary and biogeographic analyses, since such categories can then be used in connection with e.g. a phylogeny for ancestral range reconstructions (Pagel, M., Meade, A., et al. 2004, Ree, R.H. and Smith, S.A. 2008, Matzke, N.J. 2013) and for the estimation of area-dependent inferences of diversification rates (Goldberg, E.E., Lancaster, L.T., et al. 2011, Silvestro, D., Schnitzler, J., et al. 2011, FitzJohn, R.G. 2012). Species categorisation into commonly recognised areas such as eco-regions and realms (Olson, D.M., Dinerstein, E., et al. 2001, Abell, R., Thieme, M.L., et al. 2008, Holt, B.G., Lessard, J.-P., et al. 2013) may also reveal patterns of biodiversity and distribution at a large scale, facilitating the identification of regions with outstanding levels of species richness and endemism – central to the concept of Biodiversity Hotspots (Myers, N., Mittermeler, R.A., et al. 2000).

Rapidly increasing amounts of species occurrence data, the need to classify species into discrete areas in an automated, reproducible and transparent way tailored to evolutionary biologists, have led us to develop **SpeciesGeoCoder.**

## Description

SpeciesGeoCoder is written in Python (https://www.python.org/) and makes use of the R software environment (http://www.r-project.org/) for statistics and plotting of results. It runs on all platforms for which these programs are available, including Windows, MacOS X, GNU/Linux and BSD, and can be run either through the command-line or through a graphical user interface (GUI). The GUI provides researchers that are not familiar with using command-line programs, an easy-to-use set of method for analysing large amounts of geo-referenced data. The source code consists of a set of python modules, and the analyses of GeoTIFF files are done using the GDAL python bindings (http://www.gdal.org/) for fast execution. The basic workflow describing the package is illustrated in Figure 1.

**Figure 1.**
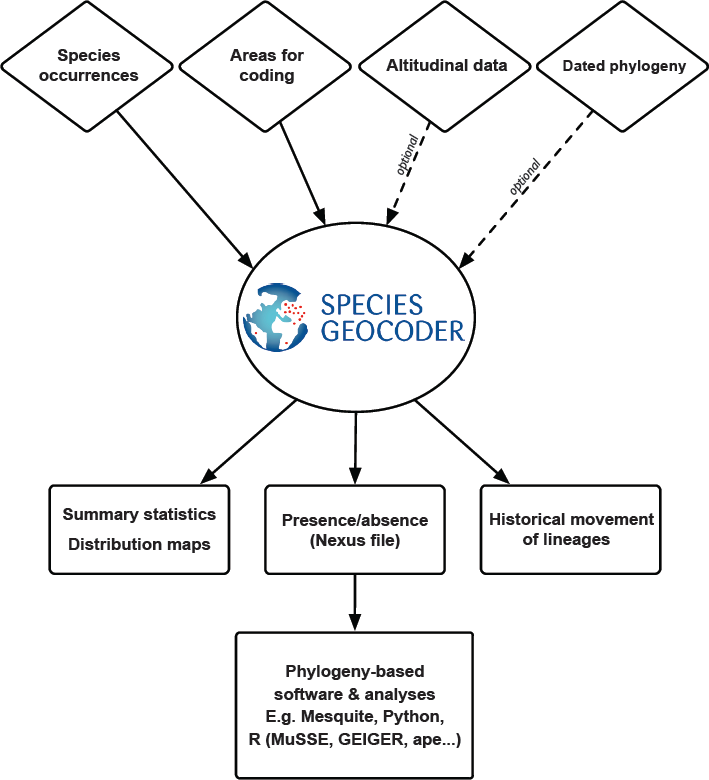
Simplified workflow of the SpeciesGeoCoder package.

SpeciesGeoCoder works as follows:

1. The user provides input data in two files: A) species occurrence data (including species names, latitude and longitude, in tab-delimited text format [see example files distributed with the program] or in the format directly provided by GBIF); and B) a text file defining the areas to be used for coding, i.e. a list of polygon names and coordinates for each of its edges, and (optionally) an altitudinal range (e.g., between 500 – 1000 meters above sea level). The polygons can be easily designed through various GIS tools (e.g. the freely available program QGIS, http://qgis.osgeo.org). If an altitudinal range is provided, additional altitudinal data files for the regions covered and at the resolution desired (in GeoTIFF format) are needed. This type of data can be freely downloaded from various on-line resources (see SpeciesGeoCoder’s wiki for a list of repositories).
2. SpeciesGeoCoder loops through all samples in the input file, counting the presences for each species in each polygon.
3. The default output is a NEXUS file (Maddison, D.R., Swofford, D.L., et al. 1997) containing a data matrix with all analysed species and their presence or absence in each area coded as ‘1’ or ‘0’, respectively. ‘Presence’ requires by default at least a single occurrence in an area, but may be set to require a number above a user-defined threshold (either an absolute number, a percentage of the occurrences, or both). Alternatively, the user may ask for a more complex output including the number of occurrences in each polygon, which could aid the identification of outliers that could require further verification; this information appears as comments in the NEXUS files. This file can be directly analysed in programmes that handle the NEXUS input format, such as Mesquite (Maddison, W.P. and Maddison, D.R. 2009) and most phylogenetic packages written in R, such as APE (Paradis, E., Claude, J., et al. 2004), GEIGER (Harmon, L.J., Weir, J.T., et al. 2008), and Diversitree (FitzJohn, R.G. 2012), as well as others written in Python, such as BayesRates (Silvestro, D., Schnitzler, J., et al. 2011).
4. The second (optional) result is a series of summary statistics and distribution maps. These include multiple PDF documents with bar charts as graphical representations of the number of species per area, the number of occurrences per species per area, and the relative occurrence per area for each species. The summary tables used for the graphical output are also provided as tab-delimited text files to facilitate downstream analyses. The distribution maps plot all occurrence points and the areas included in the analyses, coded at the species level. In addition, for datasets including less than 40 species, a co-existence matrix for each area is calculated and visualized as a heat plot.
5. The third (optional) result is a series of plots summarising the historical movements (range expansions or dispersals) of lineages between all pairs of user-defined areas, based on a dated phylogeny (or a sample of trees, to account for topological and divergence time uncertainties). These plots are computed with novel scripts that employ available packages in R, stochastic mapping of shifts in transitions along branches, and the computation of absolute as well as relative numbers of dispersals through time, i.e. corrected by the number of lineages (Silvestro, D. 2012, Fernández-Mendoza, F. and Printzen, C. 2013). Other methods for the reconstruction of ancestral areas, e.g. incorporating the possibility of vicariance and founding effects (Ree, R.H. and Smith, S.A. 2008, Matzke, N.J. 2013), may be readily used based on the output of SpeciesGeoCoder but are not implemented in the first release of this package.

## Benchmark

We evaluated the performance and scalability of SpeciesGeoCoder through a series of simulations. We focused on how computing time is determined by three key variables: *i)* the number of geo-referenced occurrences, *ii)* the number of polygons, and *iii)* the complexity of the polygons, measured by their number of edges (corners). The occurrence dataset (i) was simulated as a set of points evenly distributed longitudinally and latitudinally (see example in Fig. S1). The polygon dataset (ii) was generated in a similar way, by creating a grid of square polygons sharing two corners with each of its neighbouring polygons (Fig. S2). The edge dataset (iii) was generated by creating one square polygon and successively adding corners equally distributed over its perimeter (Fig. S3). The simulations were performed with a logarithmically increase in the number of occurrences, polygons, and polygon edges, ranging between 10^1^ and 10^8^.

Figure 2 summarises the results of these simulations, and shows that there is a linear increase in computing time in relation to the three variables examined. These results also show that SpeciesGeoCoder can handle vast amounts of data within feasible time using regular computer hardware.

**Figure 2.**
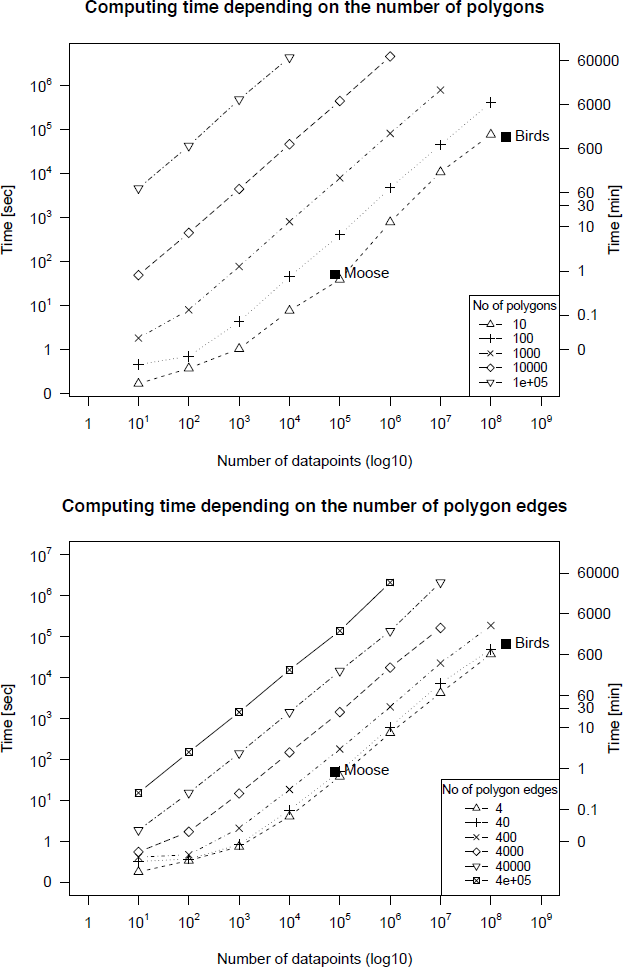
Computational time in relation to the increase in the number of polygons, polygon complexity (number of edges) and number of species occurrences, as estimated through simulations. As a comparison to empirical data, the dot denoted ‘Birds’ corresponds the coding of all c. 200,000,000 bird occurrences available from ebird.org; while ‘Moose’ corresponds to the coding of c. 80000 records, as described in the text.

## Biological Examples

### Bird biogeography

We inferred the historical dispersal of montane and lowland bird lineages through time across the Central American Seaway, which separated North and South America for millions of years until the emergence of the Isthmus of Panama. First, we downloaded the full occurrence dataset including c. 10,000 species and 200,000,000 records obtained from http://www.ebird.org (eBird 2013). We then used SpeciesGeoCoder to identify all records found outside the American continent, as well as all those found north of Mexico, which we then excluded from further analyses. Species pertaining to the remaining records were coded as occurring in Central America, South America, or both. Rather than using political boundaries, we defined the border between these two areas following the Uramita fault (Montes, C., Bayona, G., et al. 2012) that separates the South American and Panamanian geological plates (polygons shown in Fig. S4). We created two operational units from each polygon, one with the additional condition of only including records occurring below 1000 meters above sea level, and the other for records occurring above this altitudinal threshold, following the same categorisation as Weir (2006). We then reconstructed ancestral areas onto the species-level dated phylogeny of birds from Jetz et al. (2012), using stochastic mapping to reconstruct the historical dispersal of lineages through time among these four operational units. We calculated both the total (absolute) as well as the relative (in proportion to the number of lineages) number of dispersals between each pair of areas, using bins of 10 million years. Due to issues with synonymy, species concepts, and/or lack of geo-referenced data (issues which we could not readily solve computationally), the analyses dataset was reduced to include 4350 species.

Our results (Fig. 3) suggest that dispersals between the lowlands of South and Central America occurred considerably more frequently (c. 2–4 times) than dispersals between the highlands of those landmasses. There were no major differences in directionality of dispersals, except for the last time bin considered (0–10 Ma), when northwards dispersals dominated. This supports the conclusion by Weir et al. (2009) that birds mainly followed an opposite route during the Great American Biotic Interchange as compared to mammals, which migrated mostly southwards (Stehli, F.G. and Webb, S.D. 1985). The rate of dispersals increased for all categories in the most recent time bin, reflecting the full emergence of the Panama Isthmus.

**Figure 3.**
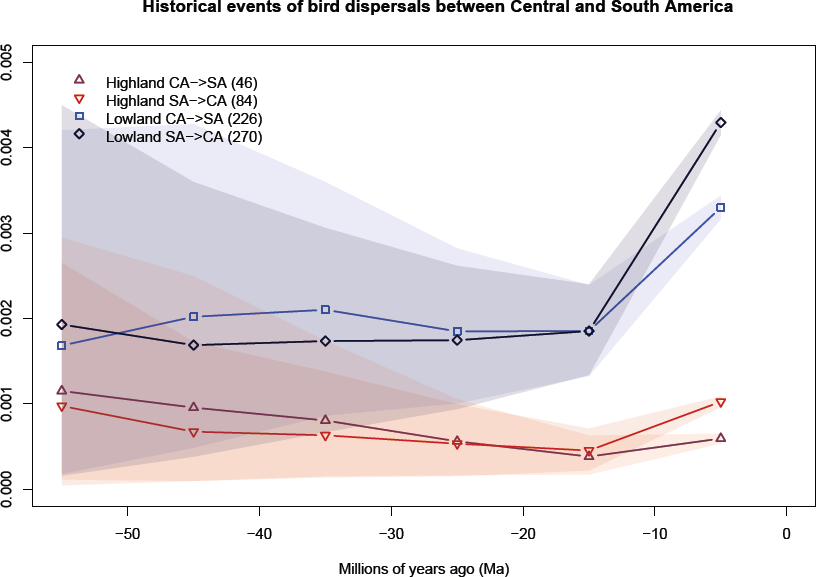
Relative frequency (Y-axis) of historical range movements (dispersals) of lowland and highland bird lineages between South America (SA) and Central America (CA), calculated per 10-million-year time bins. The total number of dispersal events inferred for our dataset is indicated between brackets in the legend.

### Seasonal behaviour in Swedish moose

We also tested the functionality of SpeciesGeoCoder at a much lower taxonomic, spatial and temporal level, by analysing a dataset of ten GPS-tracked individuals of moose (*Alces alces*) over one year in northern Sweden (c. 80,000 records). The position data were retrieved from the Wireless Remote Animal Monitoring database system for data validation and management (Dettki, H., Ericsson, G., et al. 2013). We created five arbitrary polygons, as if they would correspond to e.g. planned national parks or areas for urban development, where knowledge on wildlife activity could play a role in decision making. We calculated the co-occurrence of individuals in each of those polygons during four seasons, correcting for season length (which followed a local meteorological classification, resulting in seasons of different numbers of days: spring 60, summer 48, autumn 71, winter 186).

The results of the moose analysis are plotted together with the original data points in Fig. 4. Moose individuals spent most of the time in the north-western polygon, despite their being concentrated to a narrow strip. Co-occurrence patterns varied among seasons, with individuals being more clustered during the spring (when calves are usually born) than in the other seasons, especially in the winter (when food is more scarce).

**Figure 4.**
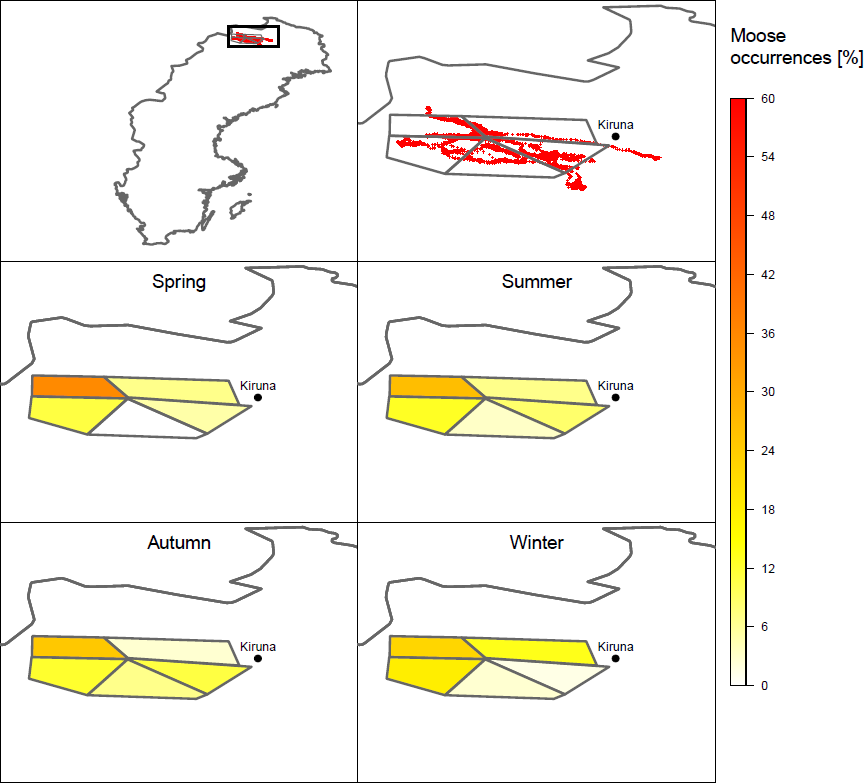
Spatial and seasonal variation of occurrences for 10 individuals of moose (*Alces alces*) in northern Sweden, scaled to the number of days in each season. The numbers on the heat scale correspond to hourly records obtained from the Wireless Remote Animal Monitoring database system and span over one year.

## Further prospects

Beyond the examples provided here, the output obtained by SpeciesGeoCoder could be readily used for e.g. calculating measures of alpha, beta and gamma diversity; the identification of neglected areas for conservation; and providing real-time detection of GPS-tagged animals entering and leaving protected areas. In addition, the visualisation and coding of species into areas may greatly facilitate cleaning up occurrence databases, by enabling the identification of outliers that may require additional examination and possible exclusion from downstream analyses.

## Availability

SpeciesGeoCoder is free software available under the GPL3 licence from https://github.com/mtop/speciesgeocoder/releases. The release and its associated Wiki page (https://github.com/mtop/speciesgeocoder/wiki) include installation instructions for Mac OSX, Gnu/Linux (and other UNIX-like systems) and Windows, bundled installation packages, example files, tutorials, a list of online repositories for GeoTIFF data, pre-defined polygons, and other useful links. A graphical user interface that provides the core functions of the program is available at http://sourceforge.net/projects/speciesgeocodergui. All pages can be accessed from http://antonelli-lab.net.

## Author contributions

A.A. and M.T. initiated the project; M.F.C. and R.S. produced the tutorial; A.Z. developed the R features; D.S. implemented the scripts for stochastic mapping; M.F.C., A.Z., R.S. and D.S. performed the simulations and analysed the biological data; M.T. led the development of the code; A.A. led the writing with contributions from all authors. All authors participated in the design and implementation of the software, the production of figures and associated material, and approved the final version of this manuscript.

## Acknowledgments

We thank Rosemeri Morokawa, Alexander Walther, students and colleagues at our labs for fruitful discussions and software testing, and Fernanda Carvalho, Henrik Nilsson and Ricardo Sawaya for comments on this manuscript. We also acknowledge Holger Dettki and Göran Ericsson for providing the moose data. Finally, we thank Paula Töpel for designing the SpeciesGeoCoder logo. Funding was provided by the Swedish Research Council (B0569601) and the European Research Council under the European Union’s Seventh Framework Programme (FP/2007-2013, ERC Grant Agreement n. 331024) to A.A., from Carl Tryggers stiftelse (CTS 11:479, CTS 12:507) to M.T., and from Fundação de Amparo à Pesquisa do Estado de São Paulo (FAPESP 2009/52161-2, 2013/10262-2) to M.F.C. The code was developed with support from the Bioinformatics Infrastructure for Life Sciences (http://www.bils.se) and tested and benchmarked on the bioinformatics computer cluster Albiorix at the Department of Biological and Environmental Sciences, University of Gothenburg.

